# Estuarine molecular bycatch as a landscape-wide biomonitoring tool

**DOI:** 10.1101/2021.01.10.426097

**Authors:** S. Mariani, L.R. Harper, R.A. Collins, C. Baillie, O.S. Wangensteen, A.D. McDevitt, M. Heddell-Cowie, M.J. Genner

**Author notes:** Correspondence to: Stefano Mariani, School of Biological & Environmental Sciences, Liverpool John Moores University, Byrom Street Campus, Liverpool L3 3AF, UK.

## Abstract

Environmental DNA analysis is rapidly transforming biodiversity monitoring and bolstering conservation applications worldwide. This approach has been assisted by the development of metabarcoding PCR primers that are suited for detection of a wide range of taxa. However, little effort has gone into exploring the value of the non-target DNA sequences that are generated in every survey, but subsequently discarded. Here we demonstrate that fish-targeted markers widely employed in aquatic biomonitoring can also detect birds and mammals present in the surrounding habitats. We showcase this feature in three temperate estuaries over multiple seasons, where dozens of bird and mammal species offer valuable insights into spatial and temporal faunal variation. Our results indicate that existing metabarcode sequence data sets are suitable for mining and exploration of this ‘molecular by-catch’, and that any future eDNA-based surveys can be designed to accommodate this enhanced property of this widely applicable tool.

## Introduction

The speed with which environmental DNA (eDNA) analysis has broadly permeated biomonitoring studies worldwide is arguably unprecedented in the history of DNA-based applications to environmental science (Hering et al. 2018; Tsuji et al. 2019). As is typically the case with novel approaches, limitations and pitfalls have led to eDNA-based methods facing much scrutiny, and technological developments remain the ongoing focus of a thriving technical literature (Deiner et al. 2017; Harper et al. 2019a; Kelly et al. 2019; Loeza-Quintana et al. 2020). However, for every caveat raised, elegant solutions are proposed, and new advantages of the methods realised (Thomas et al. 2019; Salter et al. 2019; Russo et al. 2020).

Aquatic environments have been the main beneficiaries of this ‘eDNA revolution’, largely owing to the utility of eDNA-based methods for exploring inherently poorly accessible realms, and the relative ease of collecting water, within which DNA naturally disperses, thus facilitating species detection. The utility of the methods ranges from the relatively straightforward recovery of rare (Boussarie et al. 2018) and invasive (Imamura et al. 2020) species, to more sophisticated inference on habitat gradients (Sigsgaard et al. 2019), productivity dynamics (Kelly et al. 2016; Djurhuus et al. 2020) and ecosystem structure (Aglieri et al. 2020; Harper et al. 2020). It is now possible consider aquatic eDNA as a useful tool for tackling some of the most pressing biodiversity conservation challenges in a swift, affordable and standardised way, particularly given growing interest in the generation and curation of reference DNA sequence databases. Moreover, the utility of aquatic eDNA may stretch into biomonitoring of associated terrestrial habitats. Recently, it has been shown that DNA retrieved from smaller water bodies can be used to map the distribution of terrestrial mammals that are active in proximity of the aquatic source (Harper et al. 2019b; Sales et al. 2020), suggesting water masses can act as natural biodiversity ‘collectors’.

Fundamental to the success of multi-species eDNA investigations is the choice of the genetic marker, which should be ‘universal’ across the whole taxonomic group of interest, and ‘specific’ enough to minimise the amplification of DNA from non-target taxa (Collins et al. 2019; Leese et al. 2020). As the most abundant and speciose vertebrate class on Earth, bony fishes (Osteichthyes) have played a major role in the development and consolidation of eDNA applications in marine and freshwater systems (McElroy et al. 2020), and there are now a widely recognised set of procedures that have proven successful globally (Miya et al. 2020). Interestingly, even the most efficient ‘fish’ primers tend to also amplify some DNA from other vertebrates, and whilst such components typically amount to rather pervasive biological material shed by humans and farmed animals (e.g. cattle, pig, chicken), they may sometimes unveil taxonomic records of substantial ecological and conservation value (Mariani et al. 2019).

Here we explored the concept that eDNA in estuarine areas, at the interface between land and sea, would originate from across the river drainage basin. We therefore examined samples from three UK estuaries flowing into the North Sea, collected as part of the routine monitoring operations of the UK Environment Agency, using a metabarcoding workflow designed for teleosts. Results confirm the versatility of the assay, which, beyond the 93 fish species identified as part of the primary survey, was also able to detect at least 32 birds and eight mammals, including marine, freshwater and terrestrial taxa as well as endangered and exotic species. Spatial and temporal analyses also showed significant variation in richness and community structure, which reflected the known landscape features and seasonality of the studied region. We conclude that future eDNA monitoring programmes along the coastal zone could harness this ‘molecular by-catch’ gathered by estuaries as a valuable catchment-wide biodiversity assessment tool without incurring any additional costs.

## Methods

### Data Collection

Sample locations included estuarine segments of the Rivers Tweed, Tees and Esk, situated along the North Sea coast of Britain, between 55°46’N, 1°59W and 54°29’N, 0°36W. Sites mirrored those targeted by the regular TraC survey (Environment Agency 2020), which included three netting sites each in the Tweed and Esk estuaries, and two in the Tees estuary. The Esk and Tees were surveyed in October 2016, May 2017 and October 2017, whereas the Tweed was only sampled in May and October 2017. Three 2 L water samples per site were collected immediately ahead of netting operations. Each sample was filtered through a 0.22 μm Sterivex-GP PES filter (Merck Millipore) using a 100 mL polypropylene syringe, and the filters were stored at −20°C.

We extracted DNA from filters following the mu-DNA tissue protocol (Sellers et al. 2018) and PCR-amplified an approximately 167-bp fragment of the mitochondrial 12S rRNA region using the fish-specific MiFish (Miya et al. 2015) and Teleo02 primers (Taberlet et al. 2018). Each primer pair was designed with a unique 8-bp tag to facilitate sample identification after sequencing. We then prepared three PCR-free, dual-indexed libraries using the KAPA Hyper Prep Kit, which were quantified using qPCR, pooled in equimolar concentrations, and loaded onto an Illumina MiSeq at 8pM concentration for 2×150-bp paired-end sequencing. Further details on laboratory procedures are in the Supporting Information.

Raw reads were filtered for PCR primers and demultiplexed (tag required on both ends of the amplicon, no mismatches allowed) into sample replicates using cutadapt v2.10 (Martin, 2011), followed by correction of Illumina sequencing errors (denoising) and quality filtering (default settings), using dada2 v1.16 (Callahan et al. 2016), and removal of non-homologous reads, using hmmer v3.1b2 (Eddy, 1998); further details can be found in Collins et al. (2019). Taxonomic identification followed a two-step procedure: (1) we obtained the NCBI RefSeq mitochondrion database v201 (https://www.ncbi.nlm.nih.gov/refseq/) and used the sintax algorithm in vsearch v2.15.0 (Edgar, 2016; Rognes et al. 2016) to assign a rough taxonomy; (2) we then removed reads assigned to fishes and used BLAST (https://blast.ncbi.nlm.nih.gov/Blast.cgi) to more accurately identify the remaining reads based on conditions outlined in the Supporting Information.

### Data analysis

All downstream analyses were performed in R v.3.6.3 (R Core Team, 2020). The raw data were summarised as the number of taxa and number of reads belonging to each vertebrate groups across seasons within each estuary (Fig. S1). Subsequent refinement of non-fish data, including removal of spurious taxa, correction of misassignments, false positive removal (see Fig. S2), and noise mitigation using a sequence threshold, is fully described in the Supporting Information. All fish assignments and corresponding reads were then omitted for downstream analyses.

Sequence data for PCR replicates were then pooled across biological replicates from each sampling location (Fig. S3), within each estuary. Sequence data for biological replicates taken at each sampling location were then pooled, and a bubble plot summarising eDNA detections in each estuary across different seasons produced (Fig. 1). For comparison, this bubble plot was reproduced to include species whose tissue had been sequenced in laboratories concurrently with this project and whose sequences were removed from the present data set (Fig. S4; see also Appendix 2). The pooled sequence data were converted to presence/absence using the *decostand* function in vegan v2.5-6 (Oksanen et al. 2019) for downstream analyses.

**Figure 1.**
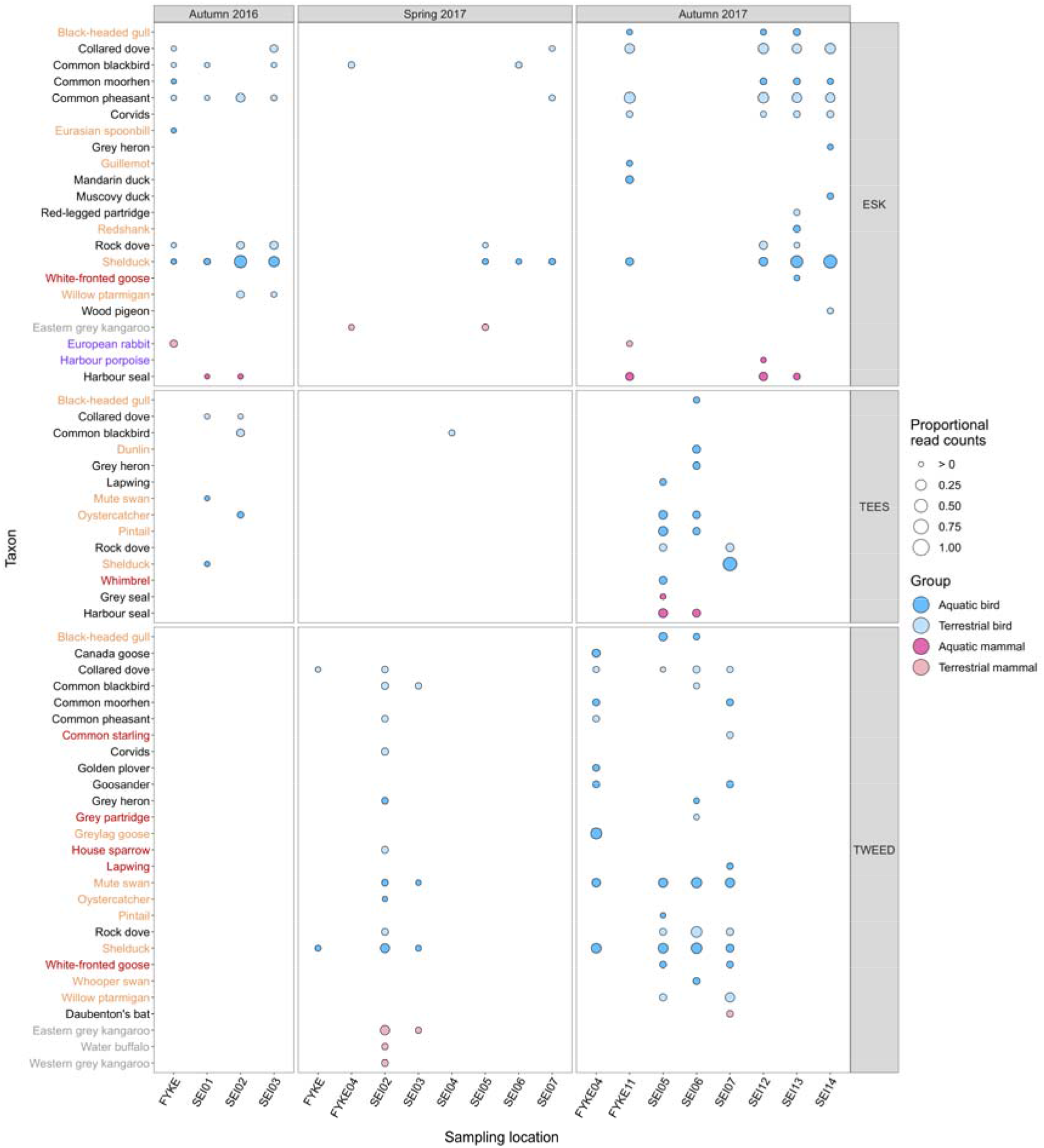
A bubble graph showing proportional read counts for taxa detected in water samples from different sampling locations within each estuary. Bubbles are coloured according to vertebrate group, and whether taxa have aquatic or terrestrial life histories. Names of birds on the Birds of Conservation Concern 4 red and amber lists (Eaton et al. 2015) are coloured red and orange respectively. Names of endangered mammals on the European Red List (IUCN 2010) are coloured purple. Names of taxa found in captivity are coloured grey.

We investigated spatial variation in α- and β-diversity between estuaries, followed by temporal variation in α- and β-diversity within each estuary, using the packages vegan v2.5-6, stats v3.6.3, FSA v0.8.30 (Ogle et al. 2020), iNEXT v2.0.20 (Hsieh et al. 2016), and betapart v1.5.1 (Baselga & Orme 2012). We define α-diversity as taxon richness of individual sampling locations, and β-diversity as the difference between communities present at each sampling location whilst accounting for taxon identity (Baselga & Orme 2012). β-diversity (Jaccard dissimilarity) was partitioned by community dissimilarity due to taxon replacement (i.e. ‘turnover’) or taxon subsets (i.e. ‘nestedness-resultant’). Details of α- and β-diversity analyses are provided in the Supporting Information.

## Results

Alongside 93 fishe species, teleost eDNA metabarcoding recovered two amphibian, 51 bird, 51 mammal, and 13 invertebrate species from 78 water samples (Fig. S1a). Most reads belonged to fishes, followed by mammals and birds (Table 1; Fig. S1b). After dataset refinement, 32 birds (21 aquatic, 11 terrrestrial) and eight mammals (three aquatic, five terrestrial) remained in 69 (88.5%) water samples. This included 18 birds and two mammals of conservation concern within Europe (Fig. 1).

**Table 1.** Summary of raw sequence output using teleost eDNA metabarcoding of estuarine water samples.

| **Group** | **Number of taxa** | **Read counts** | **Reads (%)** |
| --- | --- | --- | --- |
| Fishes | 93 | 2,379,539 | 81.563 |
| Amphibians | 2 | 88 | 0.003 |
| Birds | 51 | 178,465 | 6.117 |
| Mammals | 51 | 342,979 | 11.756 |
| Invertebrates | 13 | 16,354 | 0.561 |
| **Total** | **210** | **2,917,425** | **100.000** |

The 69 remaining samples included 33, 16, and 20 water samples from the Esk, Tees, and Tweed estuaries respectively were analysed. Alpha diversity differed across estuaries (H = 7.95, *p* = 0.018), where taxon richness was lower in the Tees than the Esk (Z = 2.263, p = 0.036) or Tweed (Z = −2.715, p = 0.020) (Fig. 2a). Taxon richness in the Esk and Tweed did not significantly differ (Z = −0.781, p = 0.435). Rarefaction and extrapolation curves indicated that lower taxon richness of the Tees may be due to differences in sample size between estuaries (Fig. 2b).

**Figure 2.**
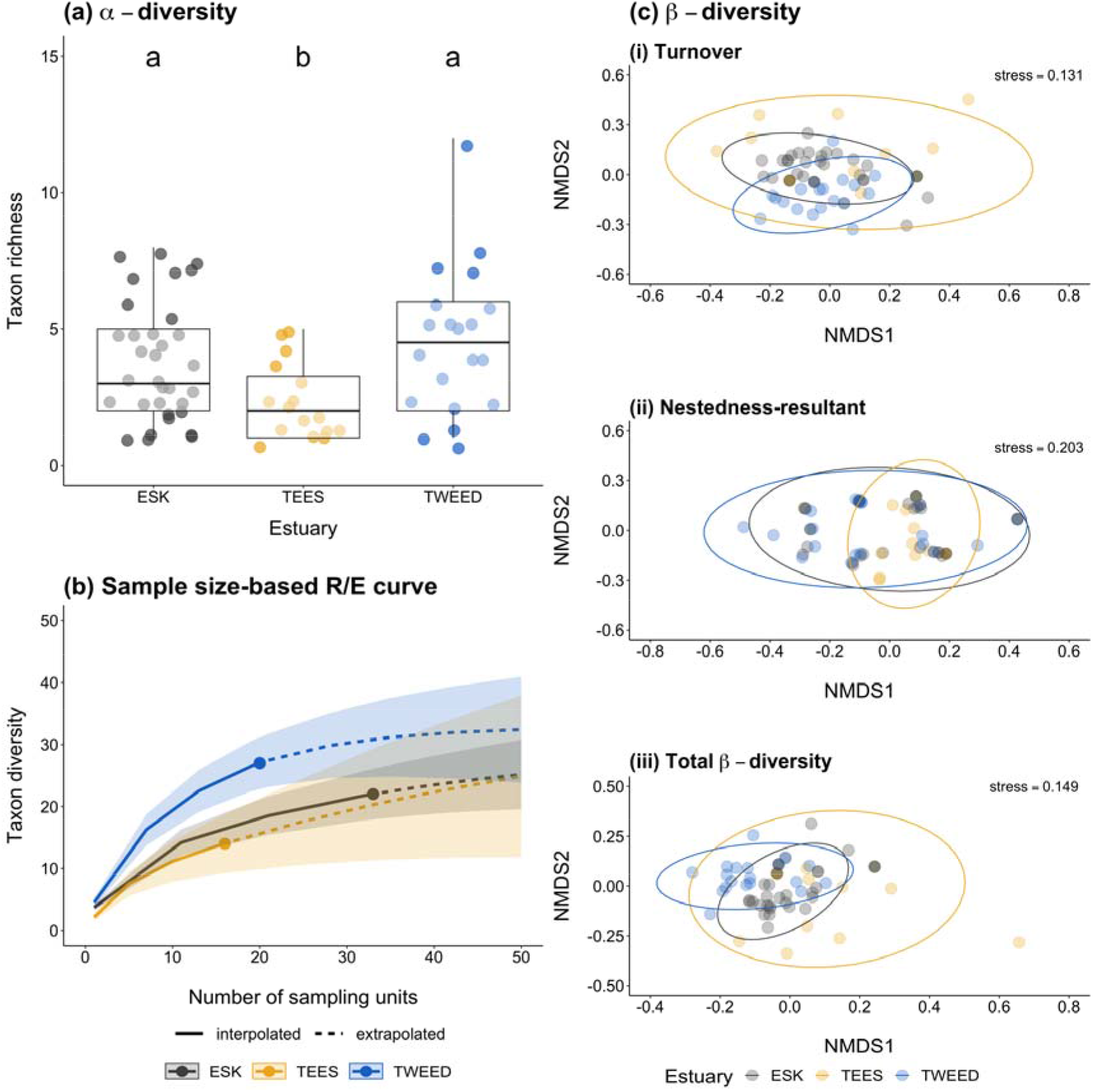
Summaries of α- and β-diversity comparisons made between sampling locations in the Esk (grey points/lines/ellipses), Tees (yellow points/lines/ellipses), and Tweed (blue points/lines/ellipses) estuaries: **(a)** boxplot showing the number of taxa detected at each estuarine sampling location, **(b)** sample size-based rarefaction/extrapolation (R/E) for each estuary, and **(c)** non-metric multidimensional scaling (NMDS) plots of estuarine communities for each β-diversity component. Letters denote significance, where different letters indicate a statistically significant difference in taxon richness derived from Dunn’s test. Boxes show 25th, 50th, and 75th percentiles, and whiskers show 5th and 95th percentiles.

Beta diversity in each estuary was driven by turnover as opposed to nestedness-resultant (Table 2). MVDISP was present between estuaries for all β-diversity components (Table 2). Estuary had a moderate positive influence on turnover (Fig. 2bi) and total β-diversity (Fig. 2biii) of communities, but not nestedness-resultant (Fig. 2bii; Table 2), generally indicating that a substantial proportion of taxa at a given estuary appear to be replaced by different taxa at other estuaries.

**Table 2.** Summary of analyses statistically comparing homogeneity of multivariate dispersions between communities at sampling locations in each estuary (ANOVA), and variation in community composition of sampling locations in each estuary (PERMANOVA). Relative contributions of taxon turnover and nestedness-resultant to total β-diversity (Jaccard dissimilarity) for each estuary are given in brackets.

|  | Homogeneity of multivariate dispersions (ANOVA) |  |  |  | Community similarity (PERMANOVA) |  |  |  |
| --- | --- | --- | --- | --- | --- | --- | --- | --- |
| | Mean distance to centroid ± SE | df | F | P | df | F | R<sup>2</sup> | P |
| *Turnover* |  | 2 | 2.822 | 0.067 | 2 | 6.014 | 0.156 | **0.001** |
| Esk (95.16%) | 0.404 ± 0.042 | | | | | | | |
| Tees (97.27%) | 0.543 ± 0.029 | | | | | | | |
| Tweed (94.21%) | 0.415 ± 0.037 | | | | | | | |
| *Nestedness-resultant* |  | 2 | 0.242 | 0.786 | 2 | 0.264 | 0.008 | 0.673 |
| Esk (4.84%) | 0.189 ± 0.026 | | | | | | | |
| Tees (2.73%) | 0.155 ± 0.017 | | | | | | | |
| Tweed (5.79%) | 0.173 ± 0.033 | | | | | | | |
| *Total β-diversity* |  | 2 | 1.839 | 0.167 | 2 | 4.203 | 0.115 | **0.001** |
| Esk (100%) | 0.522 ± 0.013 | | | | | | | |
| Tees (100%) | 0.589 ± 0.013 | | | | | | | |
| Tweed (100%) | 0.543 ± 0.011 | | | | | | | |

Alpha diversity differed across seasons within the Esk estuary (H = 20.635, *p* < 0.001) but not the Tees (H = 1.298, *p* = 0.523) or Tweed (H = 1.364, *p* = 0.243) estuaries (Fig. 3a). Taxon richness was higher in autumn (2016: Z = 2.621, p = 0.013; 2017: Z = 4.537, p < 0.001) than spring in the Esk (Fig. 3a). Furthermore, taxon richness was comparable between autumn 2016 and autumn 2017 in the Esk (Z = −1.910, p = 0.056).

**Figure 3.**
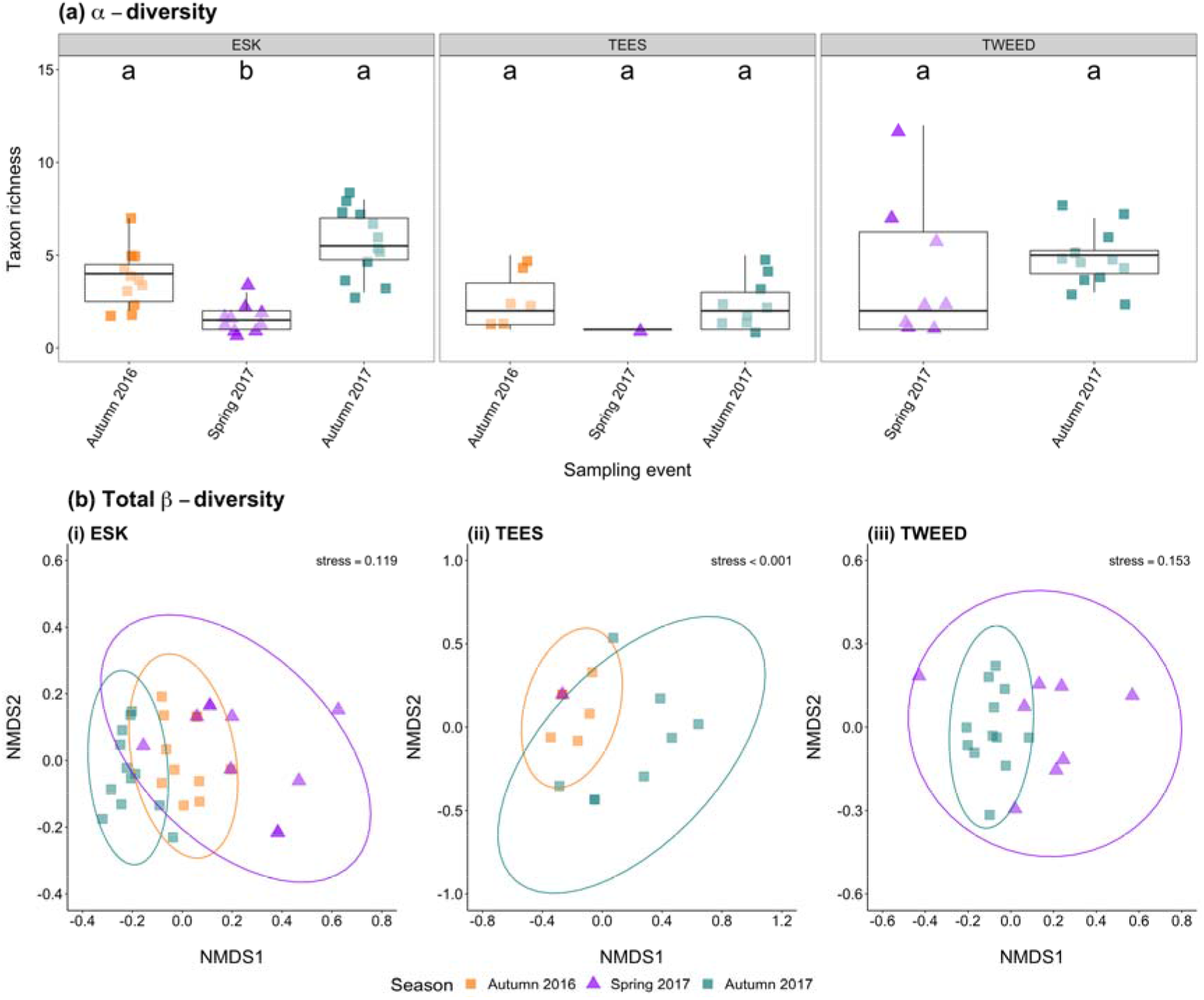
Summaries of α- and β-diversity comparisons made between sampling locations in each estuary during different seasons, including autumn 2016 (orange squares/ellipses), spring 2017 (purple triangles/ellipses), and autumn 2017 (green squares/ellipses): **(a)** boxplot showing the number of taxa detected at estuarine sampling locations across seasons, and **(b)** non-metric multidimensional scaling (NMDS) plots of communities in each estuary across seasons for each β-diversity component. Letters denote significance, where different letters indicate a statistically significant difference in taxon richness derived from Dunn’s test. Boxes show 25th, 50th, and 75th percentiles, and whiskers show 5th and 95th percentiles.

Beta diversity of estuarine communities across seasons was also largely driven by turnover, but nestedness-resultant played a greater role in some seasons. MVDISP was absent between seasons for total β-diversity (Esk), turnover and total β-diversity (Tees), and nestedness-resultant (Tweed) (Table S2). Season had a strong positive influence on all β-diversity components for the Esk (Figs. 3bi, S5ai-iii), and on turnover (Figs. S5bi, S5ci) and total β-diversity (Figs. 3bii-iii, S5biii, S5ciii) but not nestedness-resultant (Figs. S5aii, S5bii, S5cii) for the Tees and Tweed (Table S2). Therefore, taxa detected in a given season appear to be replaced by different taxa in other seasons within each estuary.

## Discussion

Since the inception of eDNA-based biodiversity assessment, there has been an emphasis on comparing detection performance with well-established biomonitoring approaches that use capture, visual or acoustic identification (Jerde et al. 2011; Foote et al. 2012; Thomsen et al. 2012; Yamamoto et al. 2016). The popularity of eDNA-based analysis today owes much to the realisation that, in many important contexts, the new tool offered significant advantages over conventional sampling methods, either through sheer improvement of detection efficacy (Boussarie et al. 2018; McElroy et al. 2020), through the discovery of its unique complementarity (Aglieri et al. 2020; Harper et al. 2020), or by simply being less resource-intensive (Bálint et al. 2018; Aglieri et al. 2020). On the other hand, little effort has gone into evaluating the intrinsically serendipitous nature of high-throughput sequencing, which, irrespective of the metabarcoding markers chosen, consistently yields substantial amounts of non-target sequences. Here we offer a demonstration that non-target sequences from metabarcoding assays contain valuable biodiversity information that can be harnessed, at no extra cost, from existing studies and ongoing surveys, dramatically expanding the reach and value of eDNA-derived data for conservation science.

We were able to conduct a multi-seasonal, parallel biodiversity survey from samples collected and analysed in three estuaries for an unrelated purpose. From a data set originally generated for monitoring coastal fish (Collins et al 2019; Siegenthaler et al 2019; Table S1), we extracted a faunal list including 32 birds and eight mammals. Of these, 52.5% were taxa that are typical of coastal marine areas, such as oystercatcher (*Haematopus ostralegus*), guillemot (*Uria algae*), common seal (*Phoca vitulina*), grey seal (*Halichoerus grypus*) and harbour porpoise (*Phocoena phocoena*). These species are directly associated with the sampled habitat, but their presence at the time of sample collection would not have been monitored by a fish-surveying team. Furthermore, some of the detected species (e.g. whimbrel (*Numenius phaeopus*), white-fronted goose (*Anser albifrons*), lapwing (*Vanellus vanellus*), redshank (*Tringa totanus*), dunlin (*Calidris alpina*), harbour porpoise) are currently listed as species of conservation concern (IUCN 2010; Eaton et al. 2015), making these DNA signatures a useful permanent record of these organisms’ presence at a certain time and space, which can serve as a baseline for future surveys, and required no financial investment to obtain.

Perhaps more surprisingly, 47.5% of the detected non-target species were not strictly associated with coastal marine areas, but rather more typical of the rural landscape, demonstrating the role of estuaries as physical collectors of eDNA transported through the drainage basin. We found ducks, passerines, waders, grouse and partridges amongst the birds, and European rabbit (*Oryctolagus cuniculus*) and Daubenton’s bat (*Myotis daubentonii*) amongst the mammals. Prior to data set refinement, a number of rodents and mustelids were also detected. Although most of these species would be expected in rural Britain, we also recovered data from species of high conservation relevance, such as the occurrence of spoonbill (*Platalea leucorodia*) in the Esk catchment, a bird that has only started breeding again in Britain in the last decade. The detection of water buffalo (*Bubalus bubalis*) as well as western and eastern kangaroo (*Macropus fuliginosus* and *M. giganteus*) in the Esk and Tweed is more puzzling. This could reflect drainage/sewage processes from nearby wildlife parks or farms: it is worth mentioning that an exotic meat company purveying both kangaroo and buffalo meat is located in the Tweed drainage, only a few miles upstream of the monitoring sites.

The utility of this ‘molecular by-catch’ in the context of landscape-wide biomonitoring is further corroborated by the marked spatial and temporal patterns identified. Taxon richness was shown to significantly vary among estuaries, and this was also reflected in the overall β-diversity configuration: the least taxon-rich estuary, the Tees, also supported a more divergent community from the other two. This can be explained by the characteristics of the catchment. Both the Tweed and the Esk run through rural landscapes, with little urbanisation, meeting the North Sea by the picturesque coastal towns of Berwick and Whitby, respectively. In contrast, the Tees flows through more urbanised areas, including the large post-industrial towns of Darlington, Middlesbrough and Hartlepool, which may arguably result in greater environmental impact on the catchment. However, rarefaction and extrapolation analyses indicated that sample coverage may have also influenced lower diversity of the Tees. With greater sample coverage, future studies may consider modelling eDNA-based results against land-use and satellite data to examine potential urbanisation and environmental gradients influencing biodiversity at landscape-scale.

The faunal records from eDNA also delineated clear temporal changes in the studied systems, with autumn samples significantly more taxon-rich than and divergent from spring samples, although this was less evident in the less diverse Tees estuary. The Esk and Tweed estuaries both supported more bird species than the Tees, including moult migrants (e.g. shelduck, *Tadorna tadorna*), winter migrants (e.g. Canada goose, *Branta canadensis;* whooper swan, *Cygnus cygnus*), passage migrants (e.g. dunlin), and partial migrants (e.g. common starling, *Sturnus vulgaris*). Additionally, more mammals were detected in the Esk and Tweed during autumn which coincides with moulting, breeding and dispersal in some species (e.g. harbour seal, *Phoca vitulina*). Autumnal influxes of birds and mammals to the Esk and Tweed may drive increased richness and community divergence, compared to less diversity in the Tees and spring generally.

The bird and mammal biodiversity ‘bonus’ showcased in this work will represent an underestimation of the actual bird and mammal eDNA diversity in the studied estuaries, and more exhaustive faunal inventories are likely to be obtained by employing taxon-specific markers for birds (Ushio et al. 2018) or mammals (Sales et al. 2020), or possibly less specific markers for vertebrates (Harper et al. 2019b). Nevertheless, the volume of information retrieved allows for educated inference on spatial and temporal variation between and within catchments, and inform and propel further focussed research activity leading to conservation actions.

In the midst of a global biodiversity crisis, rapid, powerful and affordable methods are crucial for assessing and monitoring biotas. Environmental DNA metabarcoding projects typically generate an extraordinary amount of biological information, which often exceeds the scope of the original investigation (Hupało et al. 2020). Data are routinely stored in publicly available repositories, and sequencing and computational power costs continue to drop. With this in mind, researchers and environmental managers only need to be aware of the potential of this ‘molecular by-catch’ and start designing aquatic surveys accordingly. Meanwhile, after a decade of high-throughput sequencing in natural habitats, we have already accumulated a vast amount of environmental barcodes, which remain partly untapped. We only have to start sieving through these data sets with renewed endeavour.

## Data Accessibility

Code and data to be archived in public repositories upon article acceptance.

## Acknowledgements

This study was supported by ‘*SeaDNA*’ grants NE/N005759/1 and NE/N005937/1, from the UK Natural Environment Research Council, to SM and MG. We thank Ana Soto, Naiara Sales, Riccardo Lollobrigidi, Hanna Westoby, Barry Byatt and other Environmental Agency staff for sampling support, and the *Percie Journal Club* members for offering constructive criticism on an earlier draft of the manuscript.

## Author Contributions

SM and MG conceived the study, OSW and MHC participated in sampling, OSW and CB carried out lab work, LRH and RAC analysed the data, and SM and LRH drafted the manuscript. All authors contributed to data interpretation.

